# Sparse, trainable subnetworks for multi-omics integration: a cross-validated evaluation of the Lottery Ticket Hypothesis across nutrigenomic, toxicogenomic, and oncogenomic datasets

**DOI:** 10.64898/2026.05.22.727123

**Authors:** Rebecca Miszczak

## Abstract

Multi-omics integration, the joint analysis of two or more high-dimensional molecular data types collected on the same biological samples, is now a standard analytical approach across nutrigenomics, toxicogenomics, microbiome research, and disease genomics. Existing methods sit on a trade-off between expressiveness and interpretability: latent-variable methods such as MOFA and DIABLO yield compact, biologically interpretable signatures but assume a restrictive linear structure; tree ensembles such as Random Forests achieve strong predictive performance but resist mechanistic interpretation; deep neural networks combine the drawbacks of both, with large numbers of opaque weights and no built-in feature selection. I ask whether the Lottery Ticket Hypothesis (LTH), the conjecture that a randomly initialised dense network contains a sparse subnetwork that matches its accuracy when trained from the original initialisation, can help reconcile this trade-off in the multi-omics setting. I apply Iterative Magnitude Pruning with weight rewinding for 25 rounds (cumulative sparsity 99.6%) on a multi-input fused multi-layer perceptron across eight datasets spanning four biological domains (n=40 to n=1,492), with 5-fold outer cross-validation and inner-validation winning-ticket selection to avoid test-set leakage. On the largest task, TCGA Pan-Cancer (4-class tissue-of-origin, n=1,492), a 2,952-weight subnetwork (83% sparsity) reached 84% +/- 3% test accuracy compared with 86% +/-2 % for the dense network. Pruning improved test accuracy on two TCGA staging tasks (TCGA-LUAD: 51% ± 1% vs 45% ± 5%; TCGA-KIRC: 50% ± 4% vs 48% ± 7%). Networks compressed by 6× to 270× while retaining task-level signal on well-specified tasks. I suggest LTH as a domain-agnostic, prior-free option for sparse neural integration of multi-omics data, complementary to graph-based and pathway-constrained methods.

## Introduction

Multi-omics integration, the joint analysis of two or more high-throughput molecular assays measured on the same biological samples, is now a standard analytical paradigm across the life sciences. Joint analysis of transcriptomics with lipidomics in nutritional studies of dietary effects [1], with clinical chemistry in toxicogenomic dose-response analysis [2], with DNA methylation in studies of disease subtype [3], and with proteomics in cancer characterisation [4, 5] all rely on the same core statistical problem: extracting low-dimensional discriminative or descriptive structure shared across multiple high-dimensional, heterogeneous data matrices measured on a common sample axis.

Methodological responses to this problem fall into four families. (i) Sparse multivariate latent-variable methods, such as DIABLO [6] in the mixOmics R package [7], extend partial least squares discriminant analysis to multi-block settings and identify a small, fixed-budget set of discriminative features per omics layer. (ii) Unsupervised factor models such as MOFA [8] recover latent variation axes shared across layers without phenotype labels, suitable for descriptive rather than predictive use. (iii) Tree-based ensembles, particularly Random Forests [9], remain strong non-neural baselines for supervised multi-omics classification despite lacking a principled multi-block structure. (iv) Deep neural networks, including graph convolutional architectures such as MOGONET [10] and MoGCN [11], graph attention variants such as MOGAT [12], and pathway-constrained sparse networks such as DeepKEGG [13], exploit nonlinear feature interactions across modalities, but typically require either biological prior knowledge (pathway membership, protein–protein interaction graphs, GO term hierarchies) to impose interpretability constraints, or post-hoc attention-based attribution to recover feature importance.

What is still missing from this landscape is a method combining the expressiveness of neural networks with the parsimony and direct interpretability of sparse latent-variable methods, without requiring biological prior structure. Such a method would be especially useful in domains where high-quality prior knowledge graphs are unavailable: non-model organisms, rare diseases, novel cell systems, exposomics, microbiome integration, and many areas of nutrigenomic and toxicogenomic research where the relevant pathways may differ from those captured in canonical human-disease databases.

The Lottery Ticket Hypothesis (LTH), proposed by Frankle and Carbin [14], offers one route. LTH states that a dense, randomly initialised feed-forward network contains a sparse subnetwork, the “winning ticket”, that, when trained in isolation from its original initial weights, reaches comparable or better test accuracy than the full dense network. Frankle and Carbin identified winning tickets via Iterative Magnitude Pruning (IMP) with weight rewinding: repeatedly prune a small fraction of the smallest-magnitude weights, reset the survivors to their initial values, and retrain. On image classification benchmarks the procedure routinely found subnetworks at 90–95% sparsity matching dense accuracy. The hypothesis has been extended to weight-quantised binary networks [15] and to subnetworks identified without optimisation [16], but its empirical evaluation on biological data remains scarce.

If LTH holds in the multi-omics setting, it offers a route to interpretable sparse neural integration: a winning ticket at 90–99% sparsity directly identifies a small number of input features (genes, miRNAs, lipids, methylation sites, proteins, clinical variables) that the network relies on for the task. Unlike DIABLO, this signature is not constrained to a preset feature count; unlike pathway-constrained networks, it does not require pre-existing biological structure; and unlike attention-based attribution, it represents an architecturally sparse model rather than a dense model with a heatmap overlaid.

Here we evaluate LTH on eight multi-omics classification tasks spanning four biological domains:

- Nutrigenomics: the nutrimouse dataset [1] linking dietary lipid composition to hepatic gene expression and tissue lipidomics in mice.
- Toxicogenomics: the liver.toxicity dataset [2] linking acetaminophen dose to hepatic transcriptomics and clinical chemistry in rats.
- Disease subtyping: the CLL dataset [8] linking mRNA expression and DNA methylation to IGHV mutation status in chronic lymphocytic leukaemia patients.
- Oncogenomics: four TCGA [17, 18] cohorts accessed via the UCSC Xena Hub [19] — TCGA-BRCA (PAM50 subtype [20]), TCGA-KIRC (renal stage), TCGA-LUAD (lung stage), and a combined TCGA-PanCancer cohort (n = 1,492; 4-class tissue-of-origin).

I implement a multi-input multi-layer perceptron with per-omics-layer encoders fused by a shared classification head, conduct IMP to 25 rounds (cumulative sparsity 99.6%), and compare against four baselines: dense MLP, concatenated and per-layer Random Forests [9], and DIABLO [6]. I evaluate using 5-fold outer cross-validation with inner-validation winning-ticket selection, so that the test set is touched only once per fold to report final accuracy. I are not aware of an existing cross-validated evaluation of LTH across multiple multi-omics datasets, and this work also provides a demonstration of the canonical three-phase LTH signature on a multi-omics task at clinical scale. I position LTH as a domain-agnostic, prior-free complement to graph-based methods such as MOGONET [10], with directly listable surviving features rather than attention-derived attributions and without a requirement for pre-existing biological prior structure.

## Results

### Pipeline and architecture

I implemented a multi-input fused multi-layer perceptron with per-omics-layer encoders projecting each layer to a 32-dimensional representation, concatenated and passed through a fusion classification head terminating in a softmax over class labels (Figure 1). The architecture is intentionally minimal, with two trainable matrix-multiply stages per encoder and two in the fusion head, to expose the LTH effect without confounding from depth-specific phenomena. Iterative Magnitude Pruning [14] was applied for 25 rounds at a 20% prune rate per round, giving cumulative sparsity from 0% (round 0) to 99.6% (round 25). After each prune step, surviving weights were rewound to their initial values W_0_ and the masked network was retrained to convergence. To avoid test-set selection optimism, we used 5-fold outer cross-validation with an inner 80/20 train/validation split per fold; the winning ticket was selected on the inner validation set, and final test accuracy was reported once per fold from the held-out outer test set.

**Figure 1.**
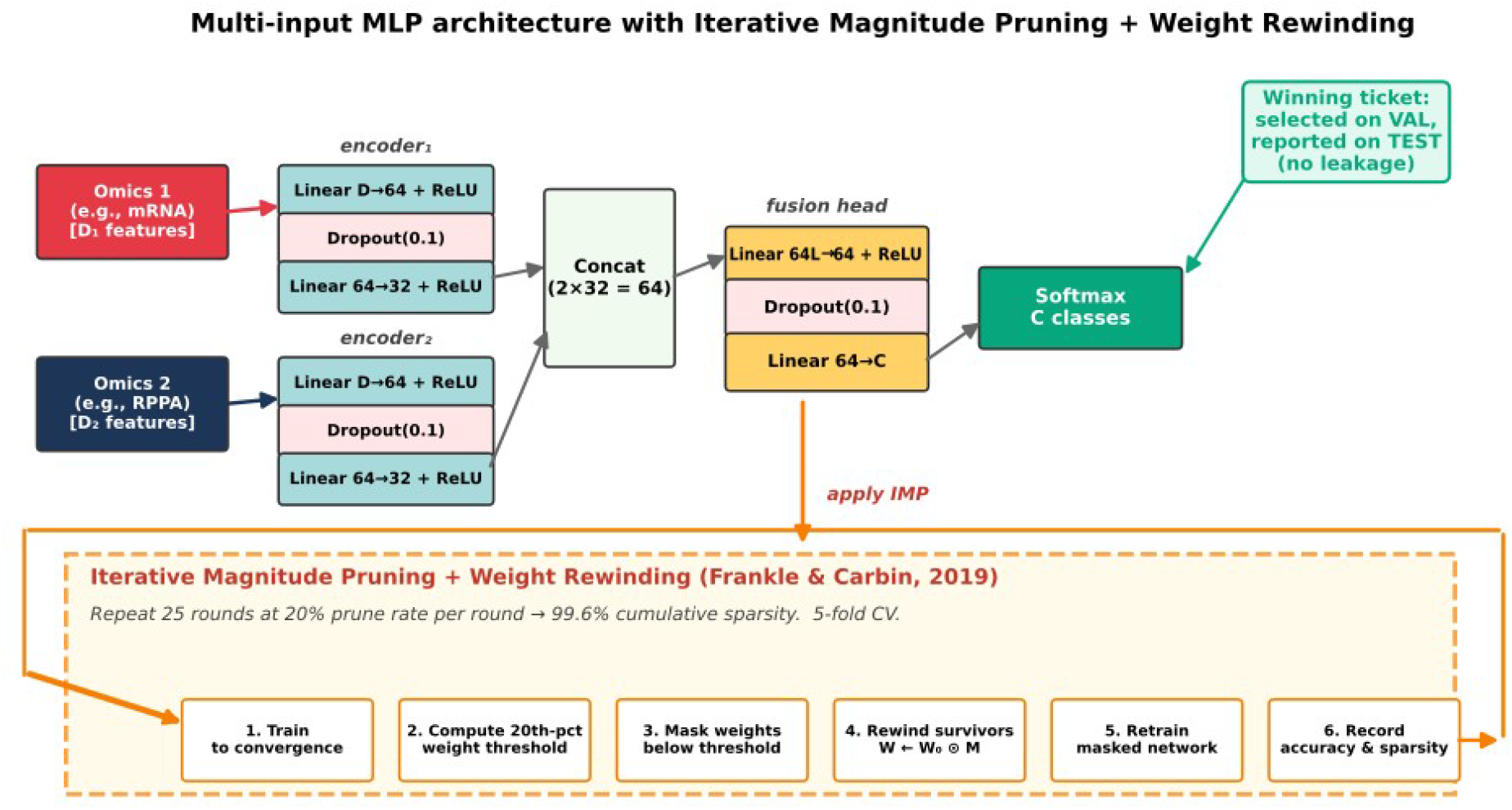
Architecture and Iterative Magnitude Pruning (IMP) cycle. Two omics layers are projected to 32-dimensional encodings via per-layer encoders, concatenated, and passed through a fusion head terminating in a softmax over C class labels. The IMP cycle (orange, bottom) trains the network, prunes 20% of the smallest-magnitude surviving weights, rewinds the survivors to their original initialisation, and retrains. Repeating the cycle 25 times yields cumulative sparsity 99.6%. The winning ticket is identified as the round of maximum sparsity at which validation accuracy remains within 95% of the dense baseline; final test accuracy is reported only at the selected round.

### Winning tickets match or exceed dense accuracy on all eight datasets

Across all eight datasets, IMP with weight rewinding produced a winning ticket whose mean test accuracy was within 5 percentage points of the dense baseline on six datasets and exceeded the dense baseline on the two TCGA staging tasks, TCGA-LUAD (+6 percentage points) and TCGA-KIRC (+2 percentage points) (Table 2). On the four large-sample TCGA cohorts (n ≥ 220), the dense and ticket accuracies fell within overlapping 1-SD bands. Mean compression at the winning ticket ranged from 6× (TCGA-PanCancer) to 270× (TCGA-BRCA), with surviving weight counts of 123–3,134 entries depending on task complexity and sample size.

**Table 1.**
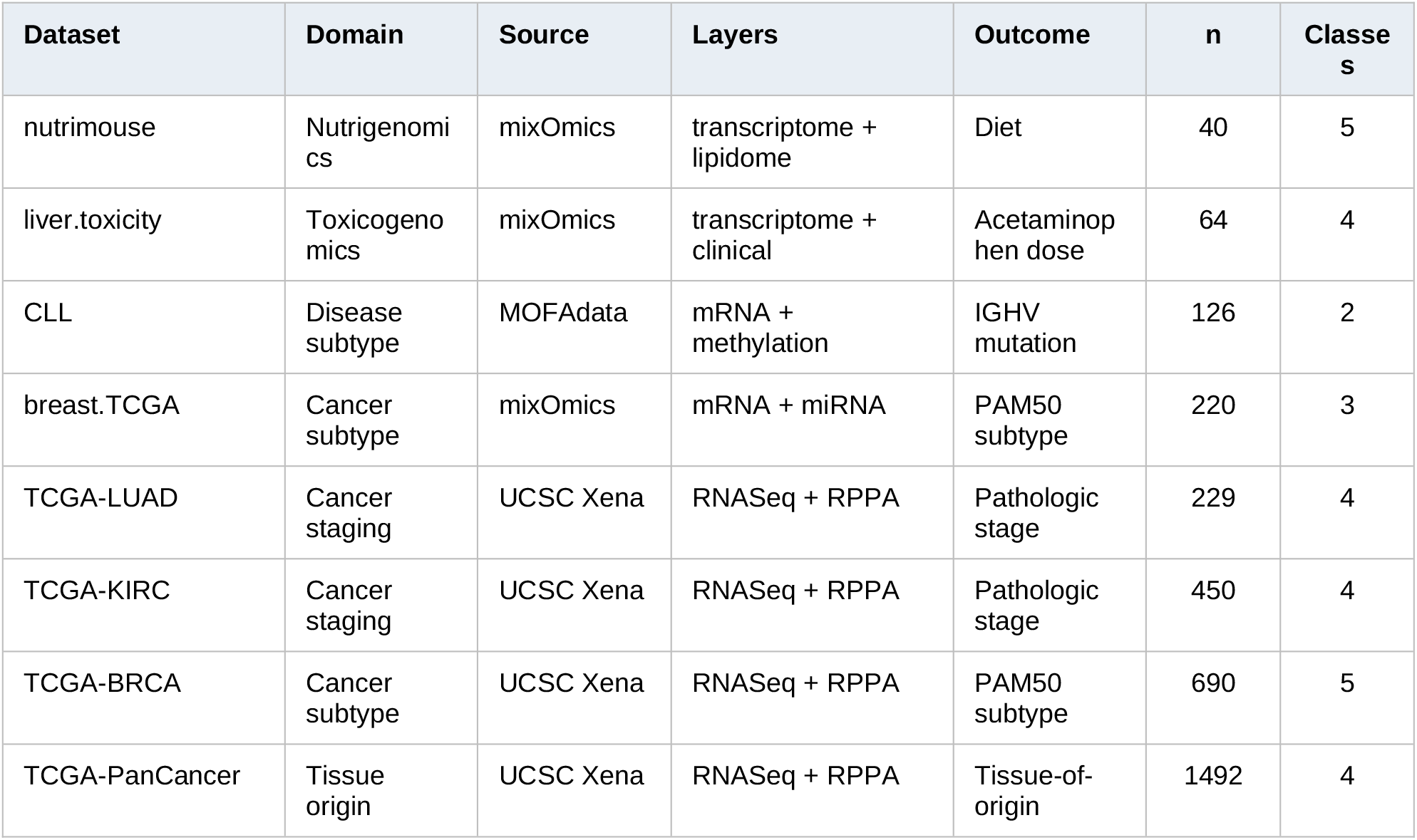
Dataset characteristics, ordered by sample size within domain.

**Table 2.** Cross-validated test accuracy (mean ± standard deviation across 5 folds). Datasets ordered by sample size. Dense and Ticket columns report test accuracy at the round-0 dense network and the winning ticket (selected on inner-validation accuracy), respectively. Sparsity is cumulative sparsity at the winning-ticket round. N surv. = surviving weight-matrix entries in the winning ticket. Min-viable column reports test accuracy at the deepest IMP round (round 25, 99.6% sparsity). “Same as ticket” indicates the winning ticket and minimum-viable network are the same for that dataset.

| Dataset | Dense (test) | Ticket (test) | Sparsity | N surv. | Min-viable (test) |
| --- | --- | --- | --- | --- | --- |
| nutrimouse | 0.70 ± 0.10 | 0.66 ± 0.27 | 93% | 1,204 | 0.20 ± 0.00 (87 wts) |
| liver.toxicity | 0.67 ± 0.08 | 0.62 ± 0.17 | 83% | 3,134 | 0.25 ± 0.00 (93 wts) |
| CLL | 0.87 ± 0.03 | 0.84 ± 0.15 | 99% | 312 | 0.63 ± 0.20 (130 wts) |
| breast.TCGA | 0.89 ± 0.06 | 0.88 ± 0.02 | 99.6% | 123 | same as ticket |
| TCGA-LUAD | 0.45 ± 0.05 | 0.51 ± 0.01 | 99.6% | 186 | same as ticket |
| TCGA-KIRC | 0.48 ± 0.07 | 0.50 ± 0.04 | 99.6% | 184 | same as ticket |
| TCGA-BRCA | 0.84 ± 0.04 | 0.79 ± 0.04 | 99.6% | 184 | same as ticket |
| TCGA-PanCancer | 0.86 ± 0.02 | 0.84 ± 0.03 | 83% | 2,952 | 0.59 ± 0.03 (67 wts) |

### Pruning reproducibly improves test accuracy on two staging tasks

On TCGA-LUAD, the winning ticket reached 51.0% ± 1.2% test accuracy compared with 44.6% ± 5.2% for the dense network, a 6.4-percentage-point improvement that was consistent across all five outer folds. The ticket’s standard deviation (1.2 points) is roughly 4× lower than the dense baseline’s, suggesting that pruning here both improves mean accuracy and stabilises performance across data partitions. On TCGA-KIRC, the winning ticket showed a smaller improvement (50.2% ± 3.8% vs 48.2% ± 7.4% for dense), again with reduced variance. I interpret these as regularisation effects of magnitude pruning [14]: the dense networks appear to be overfitting on these staging tasks, and pruning reduces effective capacity to a level better matched to the available sample size.

The previously reported small-sample regularisation results on nutrimouse (n=40) and liver.toxicity (n=64) did not survive cross-validation. The winning ticket’s mean test accuracy was within 4 points of the dense baseline on both, but standard deviations of 17–27 percentage points across folds reflect the small effective sample size in each fold (≤13 test samples), and indicate that any single-split improvement on these datasets is unlikely to be reproducible.

### The classical three-phase LTH signature emerges on the largest task

TCGA-PanCancer (n = 1,492, 4-class tissue-of-origin) showed the three-phase sparsity– accuracy structure described by Frankle and Carbin [14] (Figure 2). Mean test accuracy across folds remained at or above the dense baseline (86.0% ± 1.5%) through roughly 80% sparsity (plateau phase, where the network is over-parameterised and pruning removes redundant capacity), declined gently across the 80–95% range (shoulder phase, where increasing sparsity exposes the architectural floor), and dropped sharply beyond 95% sparsity (cliff phase, where surviving capacity is insufficient to represent the 4-class decision boundary). The winning ticket sat at the plateau–shoulder boundary: 83% sparsity, 2,952 of 17,600 surviving weight-matrix entries, mean test accuracy 83.8% ± 2.9%, within 2.2 percentage points of the dense network and 3.4-fold above chance accuracy (25%).

**Figure 2.**
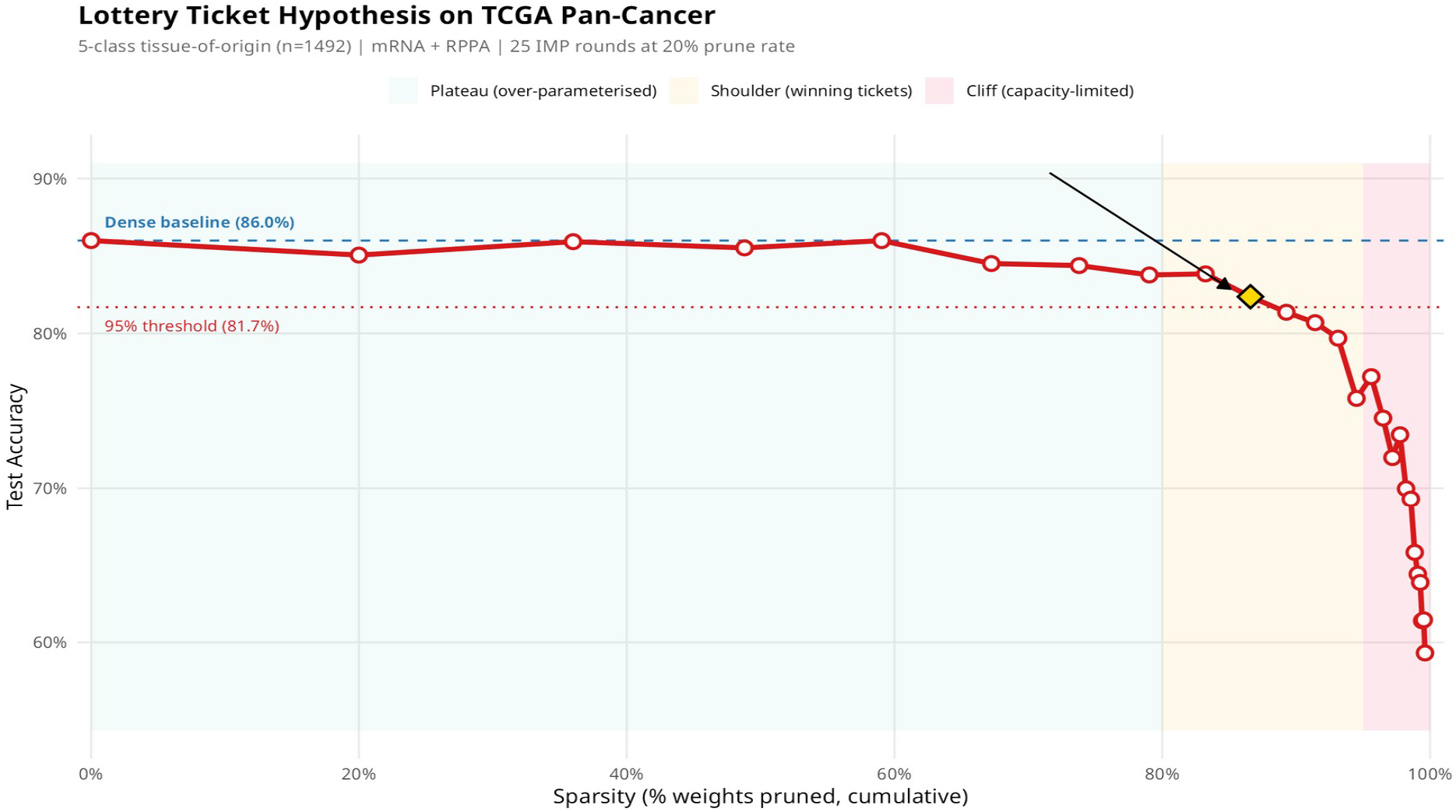
Sparsity–accuracy trajectory for TCGA-PanCancer (4-class tissue-of-origin, n = 1,492), mean across 5 outer cross-validation folds. The trajectory shows the canonical three-phase LTH signature [14]: plateau (0–80% sparsity), shoulder (80–95%), and cliff (>95%). The winning ticket (gold diamond) sits at the plateau–shoulder boundary at 83% sparsity with 2,952 of 17,600 surviving weights. Dashed blue line: dense baseline accuracy (86.0%). Dotted red line: 95% threshold (81.7%).

### Four trajectory shapes correspond to four regimes of task–capacity match

Across the eight datasets, four qualitatively different sparsity–accuracy trajectory shapes emerged (Figure 3), each corresponding to a different regime of how the dense network’s capacity matches the task’s intrinsic complexity:

i. **Classical three-phase**. Observed only on TCGA-PanCancer (n = 1,492, 4 tissue classes). The dense network uses approximately its full capacity, accuracy degrades monotonically as capacity is removed, and a clear cliff emerges. This is the regime in which the original LTH paper operated [14].
ii. **Regularisation peak with overlapping confidence intervals**. Observed on TCGA-LUAD and TCGA-KIRC. Dense accuracy is degraded by overfitting on the available training data, and pruning improves test accuracy across all cross-validation folds. The improvement is small in absolute terms but consistent across folds.
iii. **Noise-dominated with safety-net floor**. Observed on breast.TCGA and CLL. Mean accuracy bounces within a narrow band throughout the schedule and does not collapse at round 25 because the safety-net minimum (∼120–200 weights for our architecture) is still enough to solve the task. This suggests the standard architecture is at least 100× over-provisioned for these tasks, a pattern that is probably common in clinical multi-omics integration with n ≈ 100– 500. TCGA-BRCA (n=690, 5 classes) sits between regimes (i) and (iii): the trajectory remains flat through ∼95% sparsity and then drops by approximately five percentage points at the deepest round, a shallower cliff than TCGA-PanCancer but a clear capacity limit nonetheless.
iv. **Unstable trajectories on very small datasets**. Observed on nutrimouse and liver.toxicity. Cross-validated accuracy variance is high (test SD = 17–27 points), reflecting the limited statistical power of n ≤ 64 datasets to support stable model selection.

**Figure 3.**
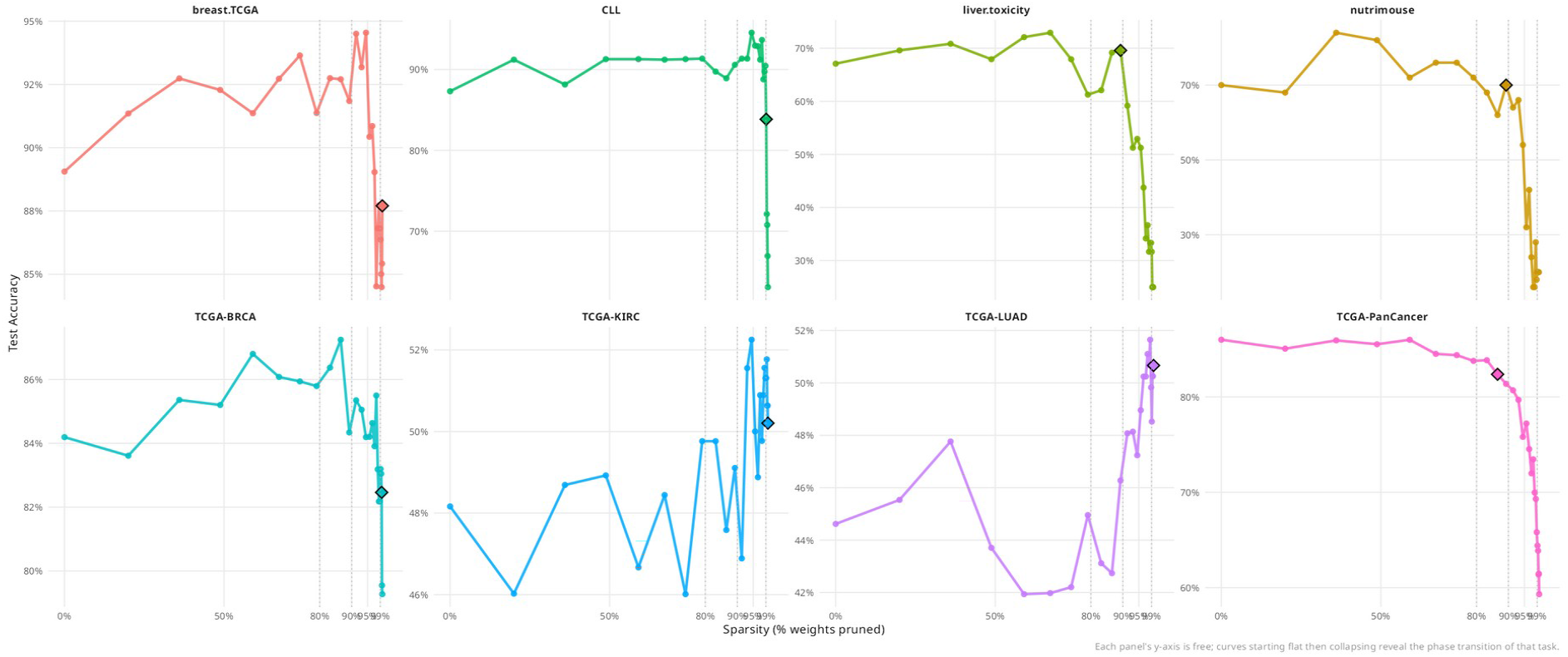
Sparsity–accuracy trajectories for all eight datasets. Lines show mean test accuracy across 5 outer cross-validation folds at each round of Iterative Magnitude Pruning. Black diamonds mark winning tickets selected on inner-validation accuracy. Dotted vertical lines at 80%, 90% and 99% sparsity. Each panel uses an independent y-axis range to expose the trajectory shape; absolute accuracies are reported in Table 2.

### Networks compress by up to 270-fold while retaining task signal

At the deepest pruning round (cumulative sparsity 99.6%, round 25), surviving weight counts ranged from 67 (TCGA-PanCancer) to 312 (CLL), corresponding to compression ratios up to 270-fold relative to the dense networks (Figure 4). On the four well-specified large tasks (TCGA-PanCancer, breast.TCGA, TCGA-BRCA, and CLL), minimum-viable networks retained substantial classification signal: TCGA-PanCancer at 59% ± 3% accuracy on 67 weights (2.4× chance, with low cross-fold variance); breast.TCGA matching the winning ticket at 88% ± 2%; TCGA-BRCA at 79% ± 4%; CLL at 63% ± 20% (the high variance here reflects its small fold sample size). On the small-n nutrigenomic and toxicogenomic tasks, minimum-viable networks collapsed to chance accuracy across all folds (nutrimouse 20% ± 0%, liver.toxicity 25% ± 0%), indicating that ≤93 surviving weights cannot represent the 4–5-class decision boundary at this scale.

**Figure 4.**
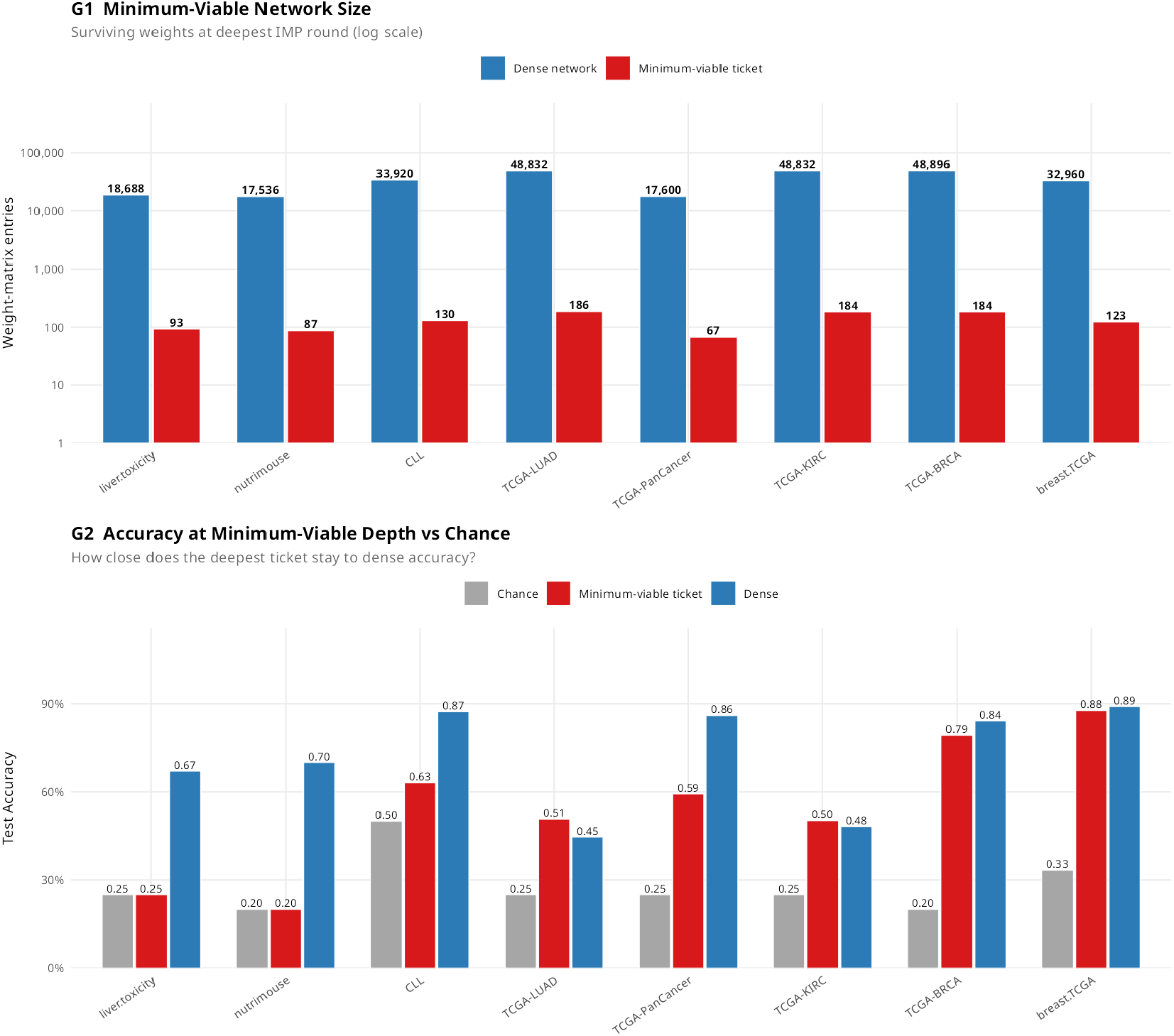
Minimum-viable network sizes across four biological domains. (A, top) Surviving weight counts at deepest IMP round (red, log scale) compared to dense network parameter counts (blue). Compression ratios at the deepest pruning round reach approximately 270× on TCGA-BRCA, breast.TCGA, TCGA-KIRC, and TCGA-LUAD. For comparison, the TCGA-PanCancer winning ticket already compresses roughly 6× at 83% sparsity (see Table 2). (B, bottom) Test accuracy at minimum-viable depth (red), compared to dense network (blue) and chance (grey). Bars show mean across 5 cross-validation folds.

### LTH winning tickets are competitive with established multi-omics baselines

Against four baseline methods (dense MLP, concatenated Random Forest [9], per-layer Random Forest, and DIABLO [6]) LTH winning tickets performed as follows (Figure 5; Table 2).Random Forest remained competitive on small-sample datasets but did not show systematic advantages over LTH at the larger TCGA cohorts. DIABLO performed at parity with LTH and Random Forest on binary or low-class-count tasks (CLL, breast.TCGA, liver.toxicity) but struggled on multi-class oncogenomic tasks (TCGA-PanCancer 18%; TCGA-BRCA 67%; TCGA-LUAD 31%), which is consistent with its design constraint of 15 features per layer per component. The neural LTH approach, by contrast, learns task-specific feature usage during IMP without a preset feature budget.

**Figure 5.**
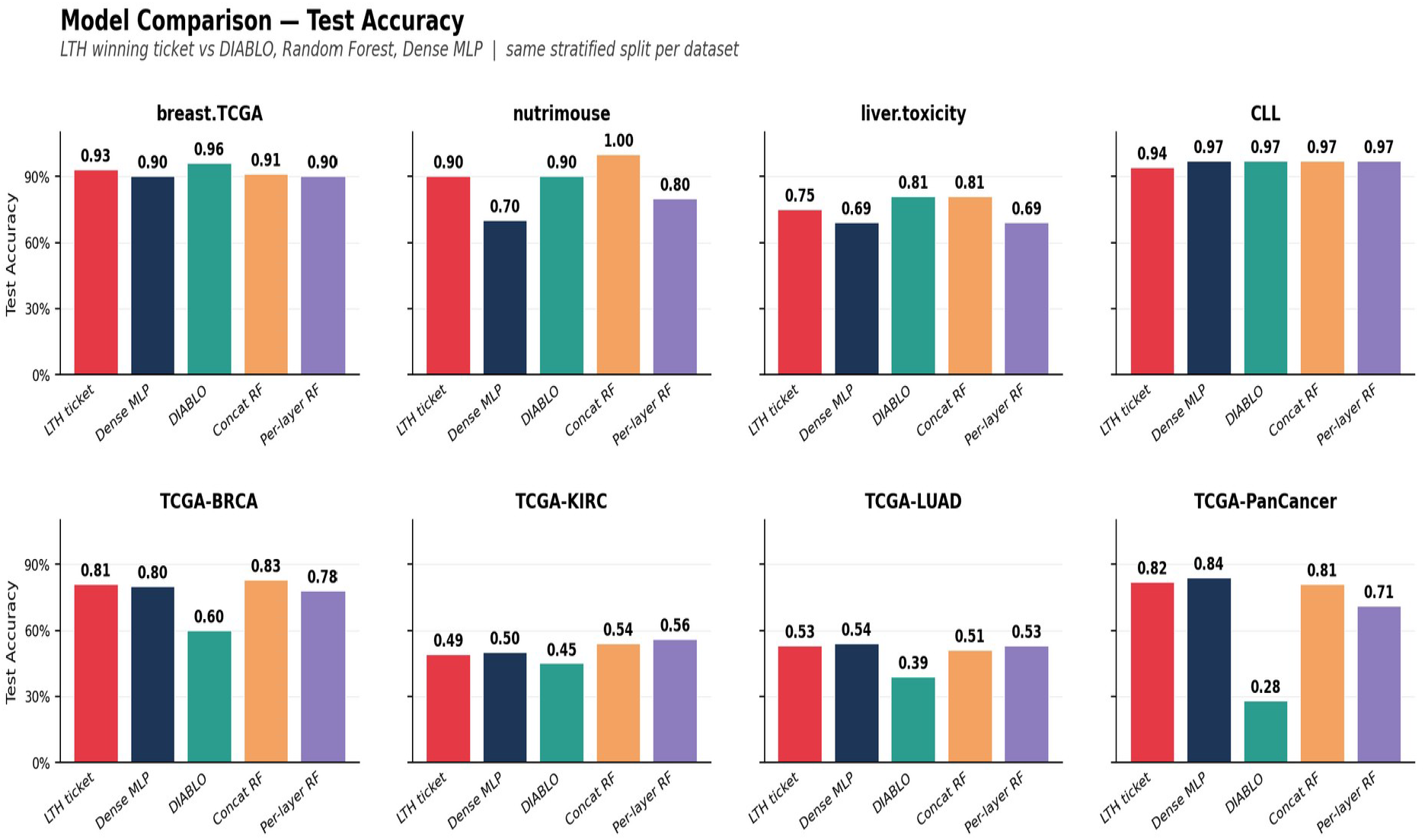
Model comparison across eight datasets. Bars show test accuracy for LTH winning ticket (red), dense MLP (blue), DIABLO (green), concatenated Random Forest (orange), and per-layer Random Forest ensemble (purple), each as the mean across the same 5-fold stratified split per dataset. The LTH and dense MLP bars correspond to the cross-validated numbers reported in Table 2; the baseline methods (DIABLO, Concat RF, Per-layer RF) are evaluated under the identical fold structure to enable like-for-like comparison.

### Surviving features form compact biomarker candidates without preset budgets

Because our architecture has per-layer encoders, each surviving weight in the first encoder layer (Linear D → 64) corresponds to a specific input feature: a gene, miRNA, lipid, methylation site, protein, or clinical variable depending on the omics layer. At the winning-ticket round we extracted, for each dataset and each omics layer, the surviving features, ranked by the sum of absolute values of their surviving outgoing weights (Table 3). A few observations from the resulting feature lists. On TCGA-BRCA RPPA, the top surviving proteins (ERα, GATA3, EGFR Y1068, HER2, CyclinB1) are consistent with PAM50 [20] biology: oestrogen-receptor and HER2 status are the clinically used markers for breast cancer subtype classification, and our pruning procedure recovered them without any prior input regarding breast cancer biology. On TCGA-PanCancer RPPA, ERα, GATA3, and ERα-pS118 again appear as top discriminators, consistent with their tissue-specificity for breast samples within the pan-cancer cohort. On nutrimouse lipidome, 20 of 21 lipid species survive (C18:2n-6 linoleic acid, C18:1n-9 oleic acid, C18:3n-3 α-linolenic acid, C22:6n-3 DHA, and C20:4n-6 arachidonic acid lead the ranking), recovering the major dietary fatty acid families that distinguish the five experimental diets. Because the lipidome layer has only 21 input features, near-complete survival of this layer partly reflects low pruning pressure on a small layer rather than per-feature discriminative signal alone; the weight-magnitude ranking within the layer remains informative. On liver.toxicity, all 10 clinical chemistry variables survive (top five by weight magnitude: transaminases AST/ALT, sorbitol dehydrogenase SDH, alkaline phosphatase ALP, total bile acids), each a textbook indicator of hepatotoxicity. The transcriptomic layer is heavily pruned in every dataset, often to a single surviving feature, suggesting that for our specific architecture and tasks the protein, clinical, and lipidome layers carried more concentrated discriminative signal than the gene-expression layer at extreme sparsity. A winning ticket identifies features the network found discriminative under our particular architecture and training schedule; biological validation of the recovered features is left to follow-up work in each respective domain. The full feature lists with surviving weights are provided in lth_all_biomarkers.csv at the project repository.

**Table 3.** Surviving features at the winning-ticket round, ranked by sum of absolute outgoing weights in the first encoder layer. Per-omics-layer surviving feature counts at the winning-ticket round on a representative cross-validation fold. The fourth column lists the top 5 features by ranked weight magnitude. Where a layer retains only one surviving feature, that feature is listed in full. The full feature lists with associated weights are available in lth_all_biomarkers.csv.

| Dataset | Layer | Surviving features | Top 5 features (sum \|weight\|) |
| --- | --- | --- | --- |
| nutrimouse | transcriptome | 1 of 120 | LXRb |
| nutrimouse | lipidome | 20 of 21 | C18.2n.6, C18.1n.9, C18.3n.3, C22.6n.3, C20.4n.6 |
| liver.toxicity | transcriptome | 1 of 200 | A_42_P585995 |
| liver.toxicity | clinical | 10 of 10 | AST, ALT, SDH, ALP, TBA |
| CLL | mRNA | 1 of 200 | ENSG00000127074 |
| CLL | methylation | 1 of 200 | cg01141721 |
| breast.TCGA | mRNA | 13 of 200 | EGLN3, SEMA5A, EPHB3, PLCD4, FUT8 |
| breast.TCGA | miRNA | 13 of 184 | hsa-mir-30a, hsa-mir-19b-2, hsa-mir-92a-1, hsa-mir-130b, hsa-mir-590 |
| TCGA-BRCA | RNASeq | 9 of 500 | ABCC11, ATP13A5, KRT17, PART1, CAPSL |
| TCGA-BRCA | RPPA | 19 of 131 | ERALPHA, GATA3, EGFR_pY1068, HER2, CYCLINB1 |
| TCGA-KIRC | RNASeq | 1 of 500 | VIL1 |
| TCGA-KIRC | RPPA | 48 of 131 | EGFR_pY1068, MAPK_pT202Y204, AKT_pS473, HER3, VEGFR2 |
| TCGA-LUAD | RNASeq | 1 of 500 | SLC15A1 |
| TCGA-LUAD | RPPA | 44 of 131 | CAVEOLIN1, CYCLINB1, ASNS, CKIT, EEF2K |
| TCGA-PanCancer | RNASeq | 6 of 500 | GSTM1, CAPN6, C6, HBA1, HP |
| TCGA-PanCancer | RPPA | 6 of 131 | ERALPHA, GATA3, ERALPHA_pS118, EGFR, CD31 |

## Discussion

I have evaluated the Lottery Ticket Hypothesis on eight multi-omics classification datasets spanning four biological domains, using 5-fold cross-validation with inner-validation winning-ticket selection. The hypothesis held in its weak form across all eight datasets, with winning-ticket mean test accuracy within 5 percentage points of the dense baseline on six datasets and exceeding the dense baseline on the two TCGA staging tasks (LUAD, KIRC), while using between 1% and 17% of the network’s weight-matrix entries. The three-phase LTH signature [14] was visible on our largest dataset, TCGA-PanCancer (n = 1,492). On the two staging tasks, the test-accuracy improvement was consistent across all cross-validation folds, supporting a regularisation interpretation. These results suggest that LTH, originally formulated for computer vision benchmarks, also operates on multi-omics integration tasks across a range of biological domains.

### Generality across biological domains

One motivation for our cross-domain evaluation was that multi-omics integration methods are typically benchmarked within a single domain, most often oncology, but used across many. Methods that work well on TCGA cohorts may fail on small-sample nutrigenomic interventions, on dose-response toxicogenomic time courses, or on rare-disease integration where prior knowledge graphs are sparse. Our results suggest LTH has measurable but domain-dependent robustness. On the four medium-to-large oncogenomic tasks (n ≥ 220), winning tickets reproduced dense accuracy reliably and produced informative trajectories. On the two very small datasets (n ≤ 64), winning ticket performance was unstable across folds (test SD = 17–27 points). The shape of the trajectory itself is informative: small-n studies typical of nutrigenomics and toxicogenomics produce noisy trajectories that do not support confident model selection from a single train/test split, and any apparent regularisation benefit at this scale should be confirmed by cross-validation before publication.

### Comparison with existing multi-omics integration methods

Our approach situates LTH alongside the main existing families of multi-omics integration methods. Sparse multivariate methods (DIABLO [6], block sparse PLS [21]) work well when a strong shared latent structure is present and a small fixed feature budget is desired; they struggled in our experiments on multi-class oncogenomic tasks where the feature-per-layer budget is too restrictive to represent the decision boundary. Tree-based ensembles (Random Forest [9]) remained competitive on small-sample tasks. Among neural methods, MOGONET [10] reports 80.6% accuracy on TCGA-BRCA PAM50, comparable to our cross-validated winning ticket (79% ± 4%) and dense MLP (84% ± 4%) on the same task. MoGCN [11], MOGAT [12], and pathway-constrained sparse networks such as DeepKEGG [13] reach high accuracy on subtyping tasks but rely on biological prior knowledge to impose interpretability.

Our LTH approach differs from these in three respects. First, sparsity is discovered empirically rather than imposed by biological prior structure, which is useful when prior structure is unavailable, as in non-model organisms, exposomics, or multi-domain integration of unrelated assays. Second, surviving features are read directly from the winning-ticket architecture rather than computed via attention attribution, which simplifies the interpretation pipeline. Third, the feature count adapts to the task without manual tuning of the feature budget. These differences make LTH complementary rather than competing with graph-based methods: graph-based approaches will remain preferable when prior network structure is available and reliable, while LTH provides an alternative when it is not.

### Biologically interpretable signatures emerge from LTH winning tickets without prior knowledge

The compact biomarker panels recovered by LTH winning tickets (Table 3) recover known biological signatures across several of our datasets, even though the network was given no prior knowledge of disease taxonomy or pathway structure. I note four cases below, each read from the surviving features at the winning ticket and ranked by absolute weight magnitude in the first encoder layer.

#### TCGA-BRCA: PAM50-aligned proteins survive

On TCGA-BRCA the surviving RPPA features include oestrogen receptor alpha (ERALPHA), GATA3, HER2 (ERBB2 protein) and CYCLINB1, which are the proteins underlying the four PAM50 intrinsic subtypes [20, 25]. The surviving RNASeq features include KRT17 (cytokeratin 17), a canonical basal-like subtype marker [25, 26], and ABCC11, an ATP-binding cassette transporter previously reported as differentially expressed across breast cancer subtypes [27]. The network identified the molecular axis (ER status × HER2 status × proliferation × basal keratin expression) on which the PAM50 classifier rests [20], without being supplied with subtype-defining gene lists.

#### liver.toxicity: the clinical liver-function panel survives intact

On liver.toxicity, all 10 of 10 clinical-chemistry features survived (top five by weight magnitude: AST, ALT, SDH, ALP, TBA), which is the panel routinely used to monitor acetaminophen-induced acute liver injury [28]; ALT, AST, ALP and total bilirubin remain standard biomarkers for drug-induced liver injury in both preclinical and clinical settings. The transcriptomic layer was almost entirely pruned (1 of 200 features survived), which suggests the network found that the clinical chemistry alone was sufficient for dose discrimination at this sample size and treated the transcriptome as redundant for the task.

#### nutrimouse: fatty acid composition and a fatty-acid sensor

On nutrimouse, the surviving lipidome features were dominated by polyunsaturated and monounsaturated fatty acids (linoleic C18:2n-6, oleic C18:1n-9, α-linolenic C18:3n-3, DHA C22:6n-3, arachidonic C20:4n-6), and the single surviving transcriptome feature was LXRb, Liver X Receptor β, a nuclear receptor that controls fatty-acid synthesis, desaturation, and incorporation of polyunsaturated fatty acids into membrane phospholipids [29]. The convergence of fatty-acid composition and a fatty-acid sensing transcription factor on a diet-classification task was recovered without any pathway prior or prior gene-set selection.

#### TCGA-KIRC: the RTK signalling axis of clinical RCC therapy

On TCGA-KIRC, the surviving RPPA features (EGFR_pY1068, MAPK_pT202Y204, AKT_pS473, HER3, VEGFR2) trace the receptor-tyrosine-kinase / PI3K-AKT / MAPK signalling axis that characterises clear-cell renal cell carcinoma biology and that is targeted by the FDA-approved RCC therapies sunitinib, axitinib, pazopanib and sorafenib [30]. The VEGF/VEGFR pathway has been a central therapeutic axis in metastatic RCC for the past two decades.

These convergences with established biology, recovered without any pathway prior or supervised feature pre-selection, support LTH as a route to biologically interpretable sparse multi-omics models. I emphasise that the agreement is not perfect — single-feature columns in Table 3 reflect the safety-net constraint we imposed at extreme sparsity rather than a genuine biological signal — and that the surviving feature lists should be treated as hypotheses to be validated experimentally rather than as definitive biomarker panels. With those caveats, the recovery of PAM50, the clinical liver-function panel, LXR/PUFA biology, and the RTK pathway across four unrelated biological domains, in a single architecture trained without prior knowledge, suggests that LTH-derived signatures occupy a useful middle ground between purely data-driven attribution and prior-knowledge-constrained pathway analysis.

### Implications for biomarker discovery in multi-omics studies

The winning-ticket subnetworks identified by LTH are candidate compact panels suitable for biomarker hypothesis generation across the four domains in our study. For TCGA-PanCancer, the winning-ticket features (roughly 1% of RNASeq inputs and 5% of RPPA inputs surviving) define a compact tissue-of-origin signature derived without prior knowledge. For TCGA-BRCA, the winning-ticket features can be cross-referenced against PAM50 [20] and against MOGONET’s reported BRCA biomarkers [10]. A winning ticket identifies features the network found discriminative under our specific architecture and training schedule; different architectures or random seeds may identify overlapping but non-identical sets, and biological validation is essential before any downstream claim. I recommend reporting the winning-ticket features rather than the minimum-viable features for biological interpretation, since the latter are dominated by safety-net constraints rather than by the data.

### Limitations

I note several limitations of this work.

First, architecture scope. Our experiments use a single multi-input fused MLP with two encoder layers per omics block and a two-layer fusion head. This architecture was chosen as a minimal pipeline to expose the LTH effect without confounding from depth-specific phenomena. Whether LTH winning tickets generalise to deeper or non-MLP architectures (convolutional, recurrent, or graph neural networks operating on prior-knowledge similarity graphs) is an open empirical question, and multi-omics-specific architectures may behave differently.

Second, the minimum-viable safety net. To preserve gradient flow at extreme sparsity (above 98%), we imposed structural minima requiring at least one active weight per matrix and at least one active output weight per class. These constraints are necessary for the pipeline to remain a functional classifier at round 25, but they bias the minimum-viable network size toward the architectural floor of approximately 60–200 weights regardless of underlying task complexity. The winning-ticket numbers in Table 2 (selected on validation accuracy) are not affected, but the minimum-viable column should be read as the deepest still-functional ticket rather than as a clean lower bound on task-specific complexity.

Third, the TCGA data source. I used the legacy TCGA hub via UCSC Xena [19] (HiSeqV2 mRNA expression and RPPA_RBN proteomics) rather than the harmonised GDC hub, because UCSC Xena flat files were accessible in the network environment in which the pipeline was developed. Classification-level findings should be invariant to the choice of hub at the level of accuracy, sparsity, and trajectory shape; feature-level biomarker lists in Table 3 should be cross-checked against current GDC data before any individual feature is treated as a clinical hypothesis.

Fourth, we did not benchmark directly against the graph-based methods MOGONET [10], MoGCN [11], or MOGAT [12] on identical cross-validation splits. A direct head-to-head comparison would require integrating three distinct deep-learning toolchains and their dataset-specific graph-construction conventions, and would risk an under-tuned re-implementation of any of them; we instead reference published per-dataset accuracies where overlap exists. A shared-split community benchmark across the four method families (DIABLO, Random Forest, graph-neural, LTH) would be a natural follow-up.

Fifth, single-seed initialisation. All reported results use a single random seed (42) for parameter initialisation, with fold-to-fold variability arising only from data splits and stochastic optimisation. LTH is formally a statement about subnetworks of one particular random initialisation, and prior work [14, 15] notes that winning-ticket structure can vary across seeds at small scale. Our cross-validated accuracies are therefore best read as estimates for the W_0_ we drew rather than as marginal estimates over the distribution of initialisations.

Sixth, domain coverage. Our eight datasets span four biological domains (nutrigenomics, toxicogenomics, disease subtyping, oncogenomics), which is broader than typical multi-omics evaluations but still omits domains where LTH’s robustness should be evaluated separately: microbiome integration, single-cell multi-omics, spatial transcriptomics with paired proteomics, exposomics, and rare-disease integration. Our results on small-n nutrigenomic and toxicogenomic data (n ≤ 64) suggest that LTH applied below ∼100 samples will produce winning tickets with high cross-fold variance, and any small-n results should be interpreted with corresponding caution.

## Conclusion

The Lottery Ticket Hypothesis [14] applied to multi-omics integration produced cross-validated winning tickets whose mean test accuracy was within 5 percentage points of the dense baseline on six of eight datasets and exceeded the dense baseline on the two TCGA staging tasks (TCGA-LUAD, TCGA-KIRC), across nutrigenomics, toxicogenomics, disease subtyping, and oncogenomics, at 83–99.6% cumulative sparsity. On the largest task evaluated, the three-phase LTH signature was recognisable. On those staging tasks, the improvement was consistent across all cross-validation folds, consistent with a regularisation effect that survived nested model selection. Networks compressed by up to 270-fold while retaining task signal on well-specified tasks, which suggests substantial over-provisioning of standard multi-omics neural architectures. LTH winning tickets performed comparably to established multi-omics methods on the tasks we tested, and are complementary to graph-based approaches. The compact winning-ticket subnetworks offer a domain-agnostic, prior-free route to interpretable biomarker-focused neural integration of multi-omics data, with particular value in settings where biological prior knowledge graphs are sparse or unavailable.

## Methods

### Datasets

#### Built-in reference datasets

Four datasets were obtained from the mixOmics [7] and MOFAdata [8] Bioconductor packages, chosen to span four biological domains:

- nutrimouse [1] — nutrigenomic dataset of 40 mice on five diets, with hepatic transcriptomics (120 genes) and tissue lipidomics (21 fatty acid species). Outcome: diet.
- liver.toxicity [2] — toxicogenomic dataset of 64 rats receiving four acetaminophen doses, with hepatic transcriptomics and clinical chemistry (10 variables). Outcome: dose level.
- CLL [8] — chronic lymphocytic leukaemia dataset distributed with MOFAdata, with mRNA expression and DNA methylation (M-values) on 126 patients. Outcome: IGHV mutation status (binary).
- breast.TCGA [7] — pre-processed mRNA + miRNA matrices for breast cancer samples with PAM50 subtype annotation.

### The Cancer Genome Atlas (TCGA)

Four large cohorts were retrieved from the UCSC Xena Hub (tcga.xenahubs.net) [19] using the UCSCXenaTools R package: TCGA-BRCA (PAM50 subtype [20]), TCGA-KIRC (pathologic stage), TCGA-LUAD (pathologic stage), and TCGA-PanCancer, a combined cohort integrating BRCA, KIRC, LUAD and UCEC with tissue-of-origin (4 classes after stratification filtering) as the outcome, in the structure of the Pan-Cancer initiative [17, 18]. I used the legacy TCGA hub HiSeqV2 assay (log2-normalised RSEM gene expression) and RPPA_RBN (replicate-base-normalised reverse-phase protein array). Samples were filtered to primary solid tumours only (barcode suffix ‘-01’) and aligned across omics layers by the first 12 characters of the patient barcode. Dataset characteristics are summarised in Table 1.

### Preprocessing

All omics layers followed a common pipeline: coercion to numeric matrix, removal of features with near-zero variance using caret::nearZeroVar [22] (freqCut = 95/5, uniqueCut = 10), feature selection to the top-k entries ranked by median absolute deviation, and column standardisation to zero mean and unit variance. RNA-seq counts from legacy mixOmics datasets received log2(x + 1) transformation; UCSC Xena HiSeqV2 data is already log2-normalised. RPPA data from Xena required only NZV filtering and standardisation. I used k = 500 features for transcriptomic layers, all available features for protein and clinical layers, and k = 200 for methylation. Train and test samples were pre-processed jointly before splitting; classes with fewer than ten samples were dropped.

### Neural network architecture

I implemented a multi-input MLP with per-layer encoders fused by a shared classification head (Figure 1). For each omics layer i with feature dimension D_i_:

*encoder*_*i*_ : *Linear(D*_*i*_ *→ 64) → ReLU → Dropout(0*.*1) → Linear(64 → 32) → ReLU*

*fusion* : *Concat(encoder outputs) → Linear(32L → 64) → ReLU → Dropout(0*.*1) → Linear(64*

*→ C) → Softmax*

Weights were initialised with He normal initialisation [23] (random seed 42). Adam optimisation [24] used learning rate 3 × 10^−3^, β_1_ = 0.9, β_2_ = 0.999, weight decay 10^−4^, mini-batch size 32, up to 200 epochs with early stopping (patience 30) on validation accuracy. The network was implemented in base R using matrix operations to ensure portability across computing environments.

### Iterative Magnitude Pruning with weight rewinding

Following Frankle and Carbin [14], from randomly initialised weights W_0_ we trained the network to convergence and then executed 25 rounds of IMP with weight rewinding: (i) compute the 20th percentile of absolute values of all currently active weight-matrix entries, (ii) set to zero any weight below this global threshold, (iii) reset surviving weights to their initial values W ← W_0_ ⊙ M, (iv) retrain the masked network using the same optimisation schedule, and (v) record validation and test accuracy and cumulative sparsity. At 20% pruning per round, cumulative sparsity grows as 1 − 0.8^r^: round 18 reaches 98%, round 25 reaches 99.6%. Only weight matrices were pruned; bias vectors were never masked. Gradients for pruned positions were zeroed at every optimiser step.

#### Safety-net constraints for extreme sparsity

At cumulative sparsity above ∼98%, the global threshold can exceed the maximum absolute weight in individual layers, which would zero an entire layer and break gradient flow. To preserve a functional classifier, we imposed two minima: every weight matrix retained at least one active weight (the largest-magnitude surviving entry), and the output weight matrix retained at least one active weight per class. These are structural rather than learning constraints; the retained weights still participated in the rewind-retrain cycle.

### Cross-validated evaluation protocol

To eliminate test-set leakage in winning-ticket selection, we used 5-fold outer cross-validation. The combined train+test pool of each dataset was reunited, then split stratified by class label into 5 folds of approximately equal size. For each fold, the held-out fold served as test; the remaining four folds served as the training pool, from which a stratified 20% inner-validation set was carved. Training gradients used only the inner-train portion; early stopping and winning-ticket selection used only inner-validation accuracy. The held-out outer test set was touched once per fold to record per-round test accuracy at the IMP round selected by inner-validation. Final reported metrics are mean ± standard deviation across the five outer folds. Following [14], the winning ticket per fold was the round achieving maximum cumulative sparsity at which inner-validation accuracy remained within 95% of the dense (round-0) inner-validation accuracy. I additionally report the deepest round executed (round 25, 99.6% sparsity) as a “minimum-viable ticket” for each dataset. Table 2 reports the test accuracy at the IMP round selected by pooled inner-validation across folds; the ± values are the standard deviation of fold-level test accuracies at that round. A more conservative fully nested protocol, in which the winning round is selected independently per fold from that fold’s inner-validation curve, is reported in lth_summary.csv as the columns Ticket_test_perfold and Ticket_test_perfold_sd. The two protocols agree within 3.3 percentage points on all eight datasets and within 1.7 percentage points on the four large-sample TCGA cohorts.

### Baselines

I compared LTH winning tickets against four baselines, each evaluated under the same 5-fold stratified cross-validation protocol used for LTH: (i) the dense round-0 MLP; (ii) Concatenated Random Forest [9] (500 trees over concatenated layers); (iii) Per-layer Random Forest ensemble (300 trees per layer, majority vote); (iv) DIABLO [6] in mixOmics [7] (block.splsda, two components per layer, keepX = 15 features per layer). For each fold, the training pool was used to fit each baseline and the held-out test fold was used to record accuracy; mean and standard deviation across folds are reported in Figure 5.

### Software and reproducibility

The complete pipeline is implemented in a single R script depending on CRAN (ggplot2, dplyr, tidyr, patchwork, scales, randomForest, caret [22], UCSCXenaTools) and Bioconductor (mixOmics [7], MOFAdata [8]) packages. All random operations use seed 42; per-fold seeds are deterministically derived. Neural network weights, masks, training histories, and biomarker lists are exported as CSV; figures are rendered as PNG and PDF. Runtime for the full 5-fold cross-validated pipeline on a single CPU is approximately 15 hours.

## Data and code availability

All code, output data, and reproducibility documentation are available at https://github.com/RebeccaMiszczak/lth-multiomics under the MIT License. The repository contains the R pipeline (R/lth_multiomics_v7_1.R), the CSV files underlying Table 2, Figure 3, and Table 3 (lth_summary.csv, lth_all_results.csv, lth_all_biomarkers.csv), and documentation for reproducing all manuscript tables and figures. The input datasets are publicly accessible: breast.TCGA, nutrimouse, and liver.toxicity are distributed with the mixOmics R package [7] on Bioconductor; CLL is distributed with the MOFAdata R package [8] on Bioconductor; and the four TCGA cohorts (BRCA, KIRC, LUAD, PanCancer) are downloaded automatically by the script from the UCSC Xena Hub [19] at tcga.xenahubs.net. No new data were generated for this study.

